# Widespread occurrence of XXY females in *Drosophila mojavensis* and related species

**DOI:** 10.64898/2026.10.21.760257

**Authors:** T. Vanderlinde, F. Uno, M. Leite, H. Bonilla, M. Vibranovski, J. Kennison, T.A. Markow, A. B. Carvalho

**Author notes:** **Corresponding authors:** Thyago Vanderlinde / A. Bernardo Carvalho, Departamento de Genética, UFRJ, CCS, Bloco A, 2o andar, sala A2-75, Ilha do Fundão, CEP 21941-617, Rio de Janeiro, Brasil, /.

## Abstract

Bridges first reported that XXY individuals are fertile females in *Drosophila melanogaster*, arising at low frequency (~1/2000) through errors in chromosome segregation (Bridges 1914; Bridges 1916). Although widely used in genetic studies, XXY females are seldom found in wild-type strains or natural populations, presumably because natural selection keeps them at very low frequency. Here we show that many wild-type strains of *D. mojavensis* and its close relatives *D. arizonae, D. navojoa*, and *D. wheeleri* carry XXY and XXYY females at frequencies of up to 88%; males from these strains are frequently XYY or XYYY. Although field collections were hampered by population declines and safety concerns, we found one XXY female (out of 87) and one XYY male (out of 36) in a natural population of *D. mojavensis*, indicating that these genotypes also occur at appreciable frequencies in nature. XXY females are expected to produce XY-bearing and X-bearing eggs in equal proportions; instead, we found a 57:43 ratio. Mathematical modeling shows that this meiotic-drive advantage is sufficient to explain high frequencies of XXY females. It is universally accepted that Y evolution responds exclusively to male selection, yet in XXY-bearing species it also responds to female fitness.

## Main

While studying the Y chromosomes in 400 *Drosophila* species (Dupim *et al*. 2018; Vanderlinde *et al*. 2025) we noticed that in four species females occasionally were PCR-positive for all Y-linked genes (henceforth, “Y-bearing females”). This differs from Y-incorporations, a type of chromosomal fusion in which females are invariably PCR-positive (Carvalho and Clark 2005; Chang and Larracuente 2017; Dupim *et al*. 2018; Chang *et al*. 2023). These four species are closely related members of the *repleta* group that live in North American deserts: *D. mojavensis, D. arizonae, D. navojoa*, and *D. wheeleri* (Oliveira *et al*. 2012; Sanchez-Flores *et al*. 2016). Careful DNA extractions and the taxonomically restricted occurrence ruled out experimental error and hinted at an undescribed phenomenon. Indeed, we found that many wild-caught strains of these species contain Y-bearing females at high frequency (up to 88%), which are absent in strains of other species of the repleta group (Table 1). Y-bearing females from *D. mojavensis* and *D. arizonae* (the best-studied species) were found in strains collected throughout their geographic ranges. Hence, Y-bearing females are not a rare abnormality but seem to be widespread within these species. We then sought to estimate their frequency in natural populations, as well as the frequency of the XYY males they produce. Unfortunately, the abundance of these species has declined drastically in recent years: two field trips in 2024 and 2025 by experienced collectors (TAM and TV) to locations previously known to harbor large populations of *D. mojavensis* and *D. arizonae* (Organ Pipe Cactus National Monument and Tucson respectively, in Arizona, USA; Las Bocas and Alamos, in Sonora, Mexico) yielded large samples of other *repleta* group species, but few *D. mojavensis*, and no *D. arizonae*. Further compounding the problem, many potential collection sites have become unsafe owing to enforcement operations against illegal immigration (US border with Mexico), or to drug trafficking (Mexican deserts). We found one XXY female (out of 87) and one XYY male (out of 36; dPCR assay) in the Organ Pipe Cactus National Monument natural population of *D. mojavensis*. We also searched for Y-bearing females in natural populations of other species of the *repleta* group from Mexico, USA, and Brazil; overall, we tested 576 wild-caught females and found none (Table S1). Hence, Y-bearing females occur at appreciable frequencies in a natural population of *D. mojavensis*, and the phenomenon seems restricted to this species and (presumably) to *D. arizonae, D. navojoa*, and *D. wheeleri*. Karyotyping ruled out translocations and chromosomal fusions and showed that Y-bearing females carry free, apparently normal Y chromosomes (Fig. 1). In the *D. mojavensis* strain 15081-1352.03, which has 88% Y-bearing females, XXYY females and XYYY males are very common, whereas in strain 15081-1352.01 XXY females appear to predominate (Table S2). The reference genome strain 15081-1352.22 (Clark *et al*. 2007) is devoid of Y-bearing females (n = 300). The high frequency of Y-bearing females in many strains of these species, despite potential fitness costs, suggests that some evolutionary force favors their spread.

**Table 1.**
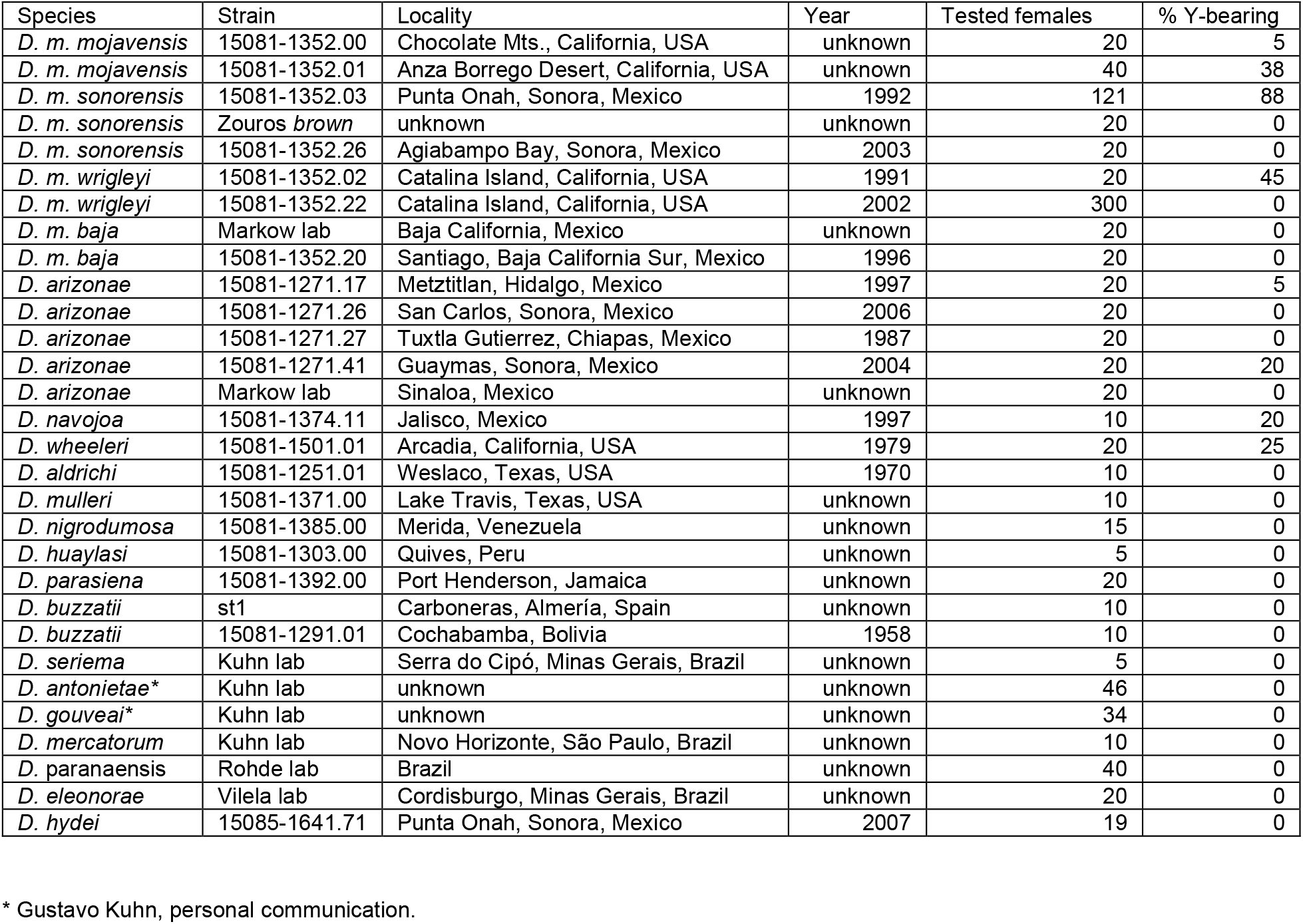
Y-bearing females in strains of *D. mojavensis* and other species of the *repleta* group.

**Figure 1.**
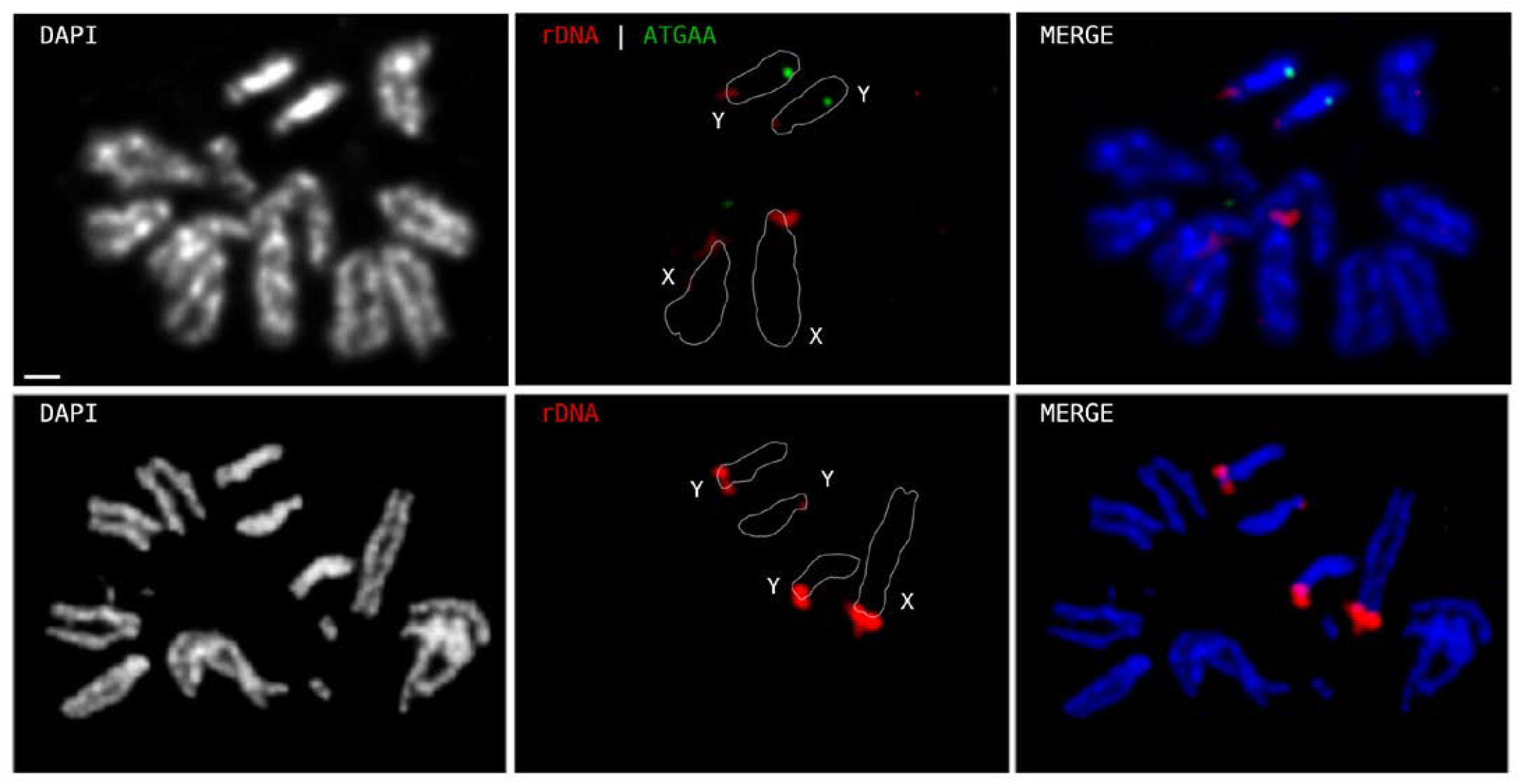
*D. mojavensis* karyotypes. Top: XXYY female, FISH staining for rDNA (red) and the Y-linked satellite ATGAA (green). Bottom: XYYY male, FISH staining for rDNA (red). All images from strain 15081-1352.03 larvae. Scale bar, 2.5 µm.

We focused the genetic analysis on *D. mojavensis*. We developed a PCR assay that distinguishes the Y chromosomes of several strains, allowing us to determine the presence and parental origin of Y chromosomes in crosses, and used dPCR to reliably quantify the number of Y chromosomes (distinguishing XY from XYY, and XXY from XXYY; Supplementary Methods; Fig. S1). Using these tools, we first characterized the transmission of the sex chromosomes in XXYY females. Their meiosis was very regular: among 162 progeny tested, 161 came from XY eggs and one from an XXY egg. This was confirmed in a larger sample genotyped with the X-linked visible marker *yellow*, which yielded an X non-disjunction rate of 0.5% (n = 804). Thus, XXYY females show regular pairing and disjunction of their two X chromosomes. Similar results were obtained in *D. virilis* (0.26% non-disjunction) (Kikkawa 1935). *D. melanogaster* XXYY females have a higher non-disjunction rate (2.9%) (Stern 1929), as do XXY females of this species (below). As expected, the daughters of the XXYY *D. mojavensis* females are XXY and carry a maternally derived Y chromosome (Table S3), whereas their sons are XYY, with one Y chromosome from each parent.

XXY females proved more interesting. They produce almost exclusively XY and X eggs: of 1,005 progeny tested, only two came from XX eggs and two from Y eggs, yielding a secondary non-disjunction rate of 0.8% (see (Cooper 1948) for the calculation of non-disjunction rate; Fig. S2; Supplementary Results). Hence, *D. mojavensis* XXY females show regular pairing and disjunction of their two X chromosomes. This contrasts sharply with *D. melanogaster* XXY females, in which the two X chromosomes frequently fail to disjoin (secondary non-disjunction: 8.2%; (Bridges 1916)), producing four egg types (XY, X, Y, and XX). Importantly, Y-bearing and XX-bearing eggs, which are frequently produced by *D. melanogaster* XXY females but not by *D. mojavensis* ones, often give rise to lethal zygotes (XXX and YY) when fertilized by sperm from normal XY males (Bridges 1916). Hence, XXY females carry an intrinsic fitness cost in *D. melanogaster*, but a much lower one in *D. mojavensis*, which probably helps to explain why they occur at appreciable frequencies only in the latter species. This difference in secondary non-disjunction rates is not unexpected. Secondary non-disjunction occurs primarily in E0 (non-exchange) tetrads (Bridges 1916; Cooper 1948; Xiang and Hawley 2006) and is therefore strongly negatively correlated with the genetic map length of the X (Sturtevant and Beadle 1936). The genetic map length of the *D. mojavensis* X chromosome (131 cM) (Staten *et al*. 2004) is more than twice that of the *D. melanogaster* X chromosome (66 cM) (Lindsley and Zimm 1992). Note, however, that the absence of meiotic costs would make the presence of Y chromosomes in females selectively neutral, but *per se* would not cause spread to high frequencies.

Unpaired chromosomes such as the Y in XXY females are expected to segregate at random; that is, XY-bearing and X-bearing eggs should be produced in equal proportions. Instead, we found that among 1,001 progeny of XXY females, 571 came from XY eggs and 430 from X eggs (57:43 ratio), a highly significant departure from the 50:50 ratio (*P* < 10^-5^, chi-square test). Such skewed meiotic segregation, known as meiotic drive, is a powerful evolutionary force that can propel even harmful traits to high frequencies (Sandler and Novitski 1957; Hartl and Clark 2007). In female meiosis, meiotic drive can result from preferential segregation of the favored allele into the functional egg, the disfavored allele going to the polar bodies (Sandler and Novitski 1957; Henikoff *et al*. 2001; Pardo-Manuel de Villena and Sapienza 2001). Hence, the widespread occurrence of XXY females in *D. mojavensis*, and their absence in *D. melanogaster* and most *Drosophila* species, appears to result from two characteristics of *D. mojavensis*: regular disjunction of the X chromosomes in XXY females (which greatly reduces the cost of lethal YY and XXX zygotes) and the meiotic-drive advantage of XY-bearing over X-bearing eggs. Both factors seem necessary: *D. virilis* XXY females (like *D. mojavensis* ones) exhibit low secondary non-disjunction (1.1%; Supplementary Text; (Demerec and Farrow 1930)), but XY eggs account for only 43% (*i*.*e*., meiotic drive favors X over XY eggs; (Kikkawa 1935)), and, accordingly, XXY females have not been reported in wild-type strains or natural populations of this species. (Bridges 1916) tested the regular daughters of *D. melanogaster* XXY females and found 80 XXY and 79 XX daughters, indicating that XY-bearing and X-bearing eggs are produced in equal proportions. *D. melanogaster* has been intensively studied for more than a century, and XXY females have been found in only one wild-caught strain, which carried a lethal X chromosome (Li *et al*. 2022) (below).

Mathematical modeling (equilibria and stability analyses; (Otto and Day 2007)) confirmed that these two characteristics of XXY females (*m*, the meiotic-drive advantage of XY eggs, and *α*, the frequency of secondary non-disjunction) are the key parameters: invasion by Y-bearing females requires *m* > *α*/2(1 − *α*), or, approximating for small *α, m* > *α*/2 (Fig. 2; this boundary assumes no selection against XYY, XXY, or XXYY genotypes; Supplementary Results). Using the experimentally derived values of *m* and *α* for *D. mojavensis*, computer simulations predict that Y-bearing females will occur at appreciable frequencies (~17%; Fig. 2), whereas for *D. virilis* and *D. melanogaster* the spread of Y-bearing females would require restrictive and probably unrealistic conditions (*e*.*g*., higher fitness of XXY females). Notably, (Li *et al*. 2022) found a *D. melanogaster* XXY strain whose X chromosomes have lost the essential ribosomal DNA (rDNA) locus, so that female survival depends on the rDNA of the Y chromosome. This case shows that, given a sufficient fitness advantage, XXY females can occur even in an unfavorable species such as *D. melanogaster*. However, it differs from that of *D. mojavensis*: it involves a lethal X chromosome that would be rapidly eliminated from natural populations.

**Figure 2.**
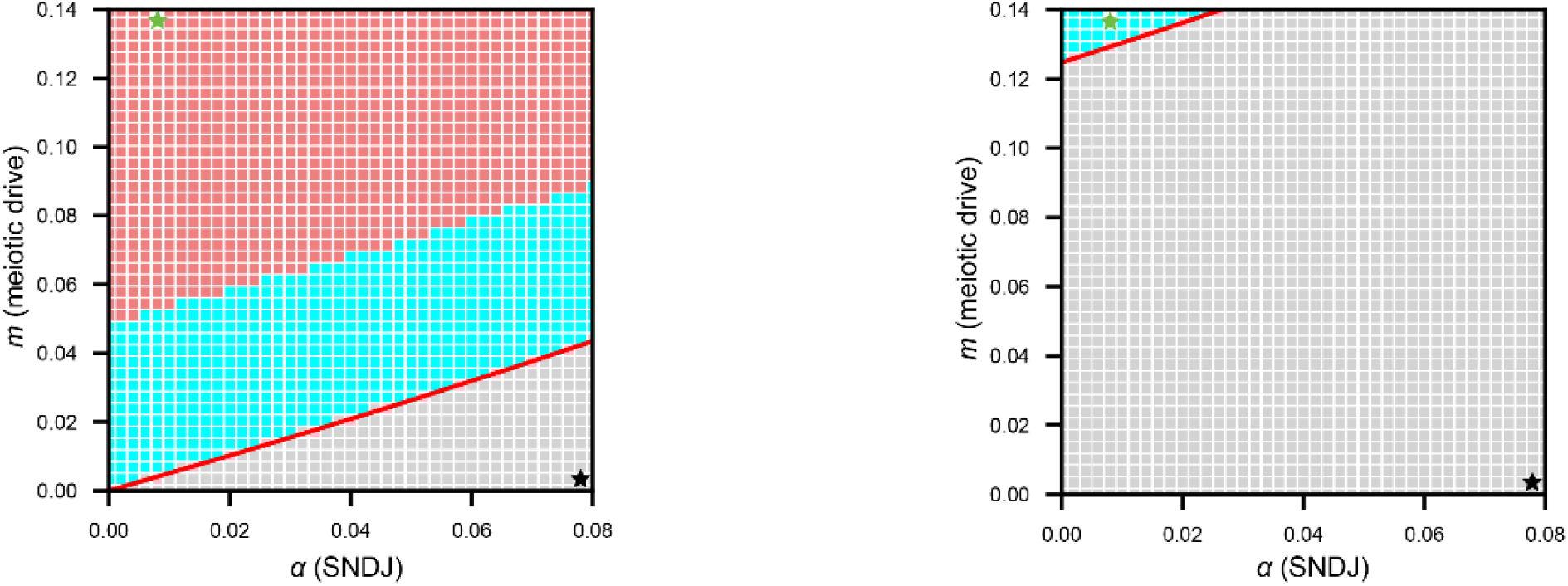
Population genetics of Y-bearing females. Mathematical modelling (red line) and computer simulations (colored squares) predict the fate of Y-bearing females, as a function of *m* (the meiotic drive of XY eggs over X eggs) and *α* (the secondary non-disjunction rate of XXY females). *m* ranges from −1 (complete meiotic drive against XY eggs) to +1 (complete meiotic drive favoring XY eggs); *m* = 0 indicates no drive (Mendelian segregation). The 57% XY eggs measured in *D. mojavensis* corresponds to *m* = 0.14. Colored squares: gray, no Y-bearing females; cyan, XXY/XX polymorphism; red, XXYY/XXY/XX polymorphism. The red line is the boundary condition (Otto and Day 2007) for invasion of Y-bearing females; note the perfect concordance with the computer simulations. Left panel: no selection against XYY, XXY, or XXYY genotypes (all fitnesses set to 1); the invasion boundary is approximately *m* > *α* / 2. Right panel: XYY fitness was set to 0.9, and the remaining genotypes to 1; note the large reduction of the parameter space that would result in Y-bearing females (see Supplementary Results for the equation of invasion boundary). The green stars in the top left of both graphs are the parameter values measured in *D. mojavensis* (*m* = 0.14; *α* = 0.008); the black stars in the bottom right are the *D. melanogaster* values (*m* = 0; *α* = 0.08). The *D. virilis* values are: *m* = −0.14; *α* = 0.011, and would be in the gray area (no Y-bearing females). The predicted frequency of *D. mojavensis* Y-bearing females (green stars) is 17.1% (left panel) and 1.7% (right).

Male parameters are also relevant because XYY males produce (and are produced by) Y-bearing females. In *D. melanogaster*, XYY males have reduced viability (~62%; (Grell 1958; Maggert 2014); Supplementary Results), whereas we found that in *D. mojavensis* the viability of XYY males does not differ significantly from that of XY males (range: 99%–109%). Mathematical modeling and computer simulations show that 62% XYY viability would block the spread of Y-bearing females, even under the favorable *m* and *α* values of *D. mojavensis*; even 90% XYY viability would strongly reduce the likelihood of finding Y-bearing females (Fig. 2). Essentially the same results were obtained by applying selection against XXY females (Supplementary Results). This sensitivity to relatively minor fitness differences offers a plausible explanation for the large difference in the frequency of Y-bearing females between laboratory strains and natural populations. We also investigated the meiotic segregation of XYY males in *D. mojavensis* and found that they produce four types of sperm: X, YY, XY, and Y (30%, 29%, 24%, and 17%, respectively; n = 358). The same four sperm types were produced by *D. melanogaster* XYY males, but at different frequencies (11%, 7%, 41%, and 41%, respectively; (Grell 1958)). Mathematical modeling and computer simulations showed that, without selection against XYY males, these male meiotic parameters do not affect the *m* > *α*/2 boundary for invasion by Y-bearing females, although they do change the relative amounts of XXYY and XXY females at equilibrium (Fig. S3; Supplementary Results). With selection against XYY males, the invasion boundary shifts and its equation becomes more complex (Supplementary Results).

Although we focused here mainly on meiotic effects, Y chromosomes can affect females in other ways. (Cooper 1956) noted that *D. melanogaster* XXYY females are “crippled”, showing leg abnormalities and reduced fecundity; careful examination of *D. mojavensis* XXYY and XXY females revealed no morphological abnormalities. The *D. mojavensis* crosses designed to measure meiotic parameters of XXYY and XXY females also produced XX females, allowing us to compare their fecundities; we found no significant difference, although statistical power was low (Supplementary Results). The few studies on the fitness of XXY females in *D. melanogaster* used attached-X chromosomes, which introduce serious bias (*e*.*g*., XXY advantage may reflect the Y chromosome rescuing rDNA-poor attached-X chromosomes; Supplementary Discussion). Y chromosomes can also affect females through the expression of Y-linked genes. RNA-seq experiments on *D. mojavensis* XXYY and XXY larvae showed that they do not express Y-linked protein-coding genes (Supplementary Results). Using the same data, we found that XXY (and probably XXYY) females do express the Y-linked copies of rDNA, as observed in *D. melanogaster* (Warsinger-Pepe *et al*. 2020). Y chromosomes are also known to have widespread effects on gene expression in males and XXY females of *D. melanogaster* (Lemos *et al*. 2008; Lemos *et al*. 2010; Elgin and Reuter 2013), and the same probably holds for *D. mojavensis*. The relevance of these findings, if any, to the Y-bearing female phenomenon remains to be determined. In any case, female meiotic parameters alone provide a compelling explanation for the presence of Y-bearing females in *D. mojavensis*, and their absence in *D. melanogaster* and *D. virilis*.

Should we expect to find other cases of Y-bearing females? The trait can evolve only in species in which, as in *Drosophila*, the primary sex-determination signal is not the presence of a Y chromosome (or a W). This information is unavailable for most species with sex chromosomes, but known cases include chicken, some fishes, *Drosophila*, and several plants (Ainsworth *et al*. 1998; Bachtrog *et al*. 2014). We should add that, although we examined 400 *Drosophila* species (Dupim *et al*. 2018), for the vast majority we sampled only a few individuals from a single strain. Finally, it is worth noting that *D. mojavensis* and *D. arizonae* are well-studied models of speciation and ecology (Vigneault and Zouros 1986; Krebs and Markow 1989; Clark *et al*. 2007; Allan and Matzkin 2019), and yet Y-bearing females had escaped detection. Hence, it is not unlikely that other cases will be found.

How did the XXY system evolve? An X chromosome with low secondary non-disjunction is certainly a facilitating factor, as is normal viability of XYY males, since both factors lower the cost of Y-bearing females. However, some direct advantage such as meiotic drive of XY over X eggs is required. As *D. virilis* and *D. melanogaster* show, meiotic drive favoring XY eggs is not an automatic consequence of meiosis mechanics in XXY females. Instead, its occurrence in *D. mojavensis* plausibly resulted from the same evolutionary mechanism that leads to conventional (male) Y-linked meiotic drive: Y chromosomes that increase their own transmission will spread in populations (Hamilton 1967). The striking difference is that in *D. mojavensis* this would have been achieved through female meiosis. A possible scenario for the evolution of Y-bearing females in *D. mojavensis* and related species is outlined in Fig. 3: a Y chromosome without meiotic drive (*i*.*e*., XXY females producing XY- and X-bearing eggs in equal proportions) is predicted to be replaced by a new variant with weak meiotic drive, in which XXY females produce 52% XY eggs. The absence of major costs would have allowed the weakly driving Y to attain a high frequency, facilitating the gradual strengthening of meiotic drive through the accumulation of new Y-linked mutations and enhancers. The end result may be either fixation of the stronger driver (Fig. 3), or a balanced polymorphism maintained by some deleterious effect of the driver (Carvalho *et al*. 1997). The absence of Y-bearing females in some *D. mojavensis* strains (Table 1) may be due to a non-driving Y; we are currently investigating this possibility. Whatever the origin of the system, the current dynamics of Y chromosomes in *D. mojavensis* and related species are influenced by meiotic drive in XXY females. Hence, while it is universally accepted that Y chromosome evolution responds exclusively to selection in males, in these species the fate of Y-linked alleles is also determined by their effects in females.

**Figure 3.**
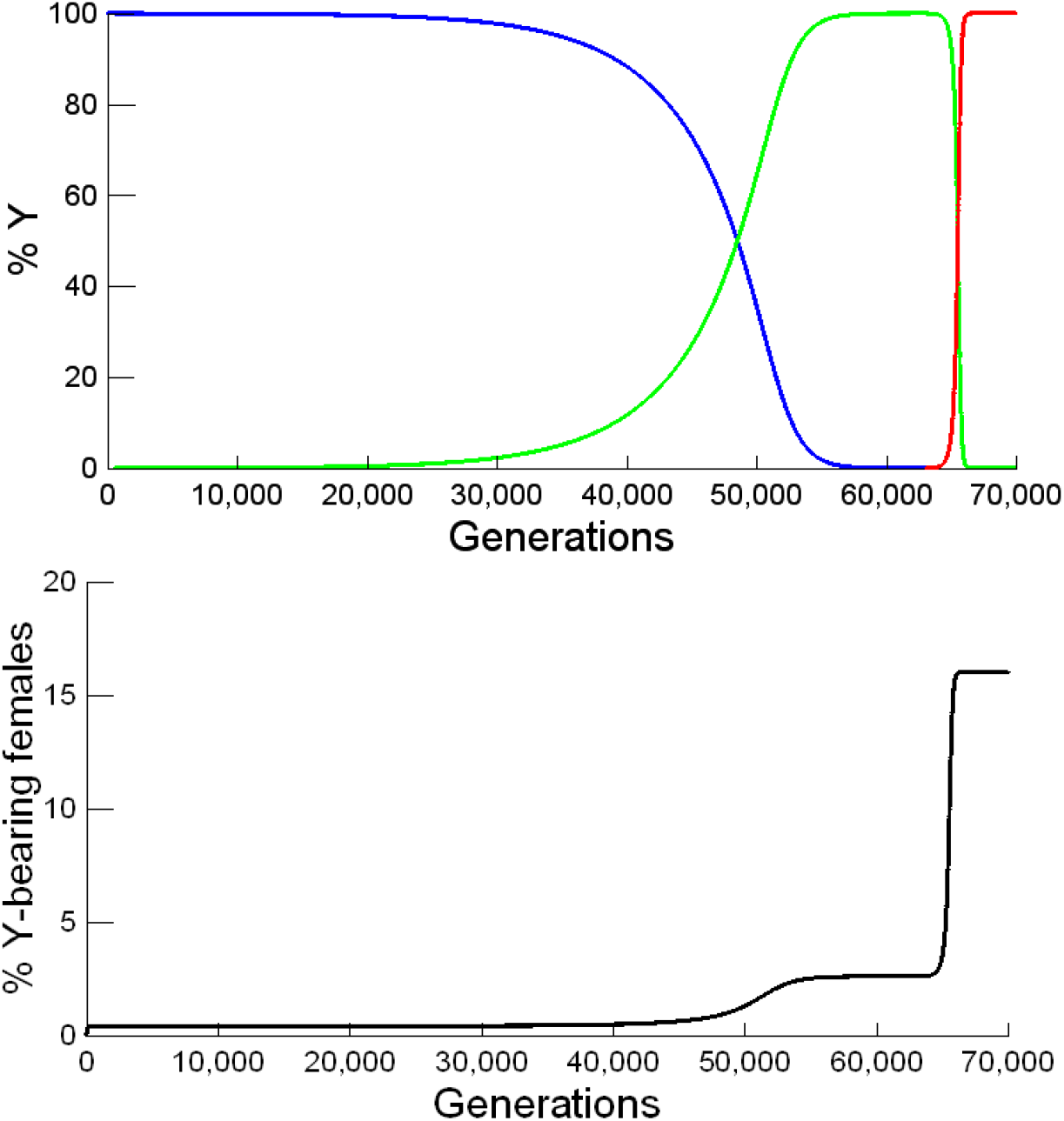
Dynamics of Y-bearing females. Top: frequency of Y chromosome variants; bottom: percent Y-bearing females. The simulated population started with only XX females and XY males, and a non-driving Y chromosome (blue line). Around generation 700, the population reached an equilibrium between primary and secondary non-disjunction, with Y-bearing females at a frequency of 0.4% (black line). Once equilibrium was reached, a weak Y driver (52% XY eggs; *m* = 0.04; green line) was introduced at low frequency (1/10,000); it gradually replaced the non-driver Y, increasing the frequency of Y-bearing females to 2.6%. Finally, a stronger Y driver (57% XY eggs; *m* = 0.14; red line) was introduced at low frequency and quickly replaced the weaker one, increasing the frequency of Y-bearing females to 16%. Male and female meiotic parameters were set to the estimated values for *D. mojavensis* (see text). Given the lack of a primary non-disjunction estimate for *D. mojavensis* we used the *D. hydei* value of 1/13,000 (Gregg and Day 1965). We assumed very weak selection (1%) against XYY, XXY, and XXYY genotypes.

## Acknowledgements

We thank Suzana Vaz and Gustavo Kuhn for valuable suggestions during this work. This research was funded by the Wellcome Trust, grant 207486Z17Z; FAPERJ–Fundação Carlos Chagas Filho de Amparo à Pesquisa do Estado do Rio de Janeiro, grant CNE2018; CNPq–Conselho Nacional de Desenvolvimento Científico e Tecnológico, grant INCT-EM, to A.B.C. F.U. is supported by CAPES–Coordenação de Aperfeiçoamento de Pessoal de Nível Superior, Finance Code 001.

